# The establishment of locally adaptive inversions in structured populations

**DOI:** 10.1101/2022.12.05.519181

**Authors:** Carl Mackintosh, Michael F Scott, Max Reuter, Andrew Pomiankowski

**Affiliations:** Department of Genetics, Evolution, and Environment, University College London, Gower Street, London, WC1E 6BT; CoMPLEX, University College London, Gower Street, London, WC1E 6BT; School of Biological Sciences, University of East Anglia, Norwich Research Park, Norwich, NR4 7TJ

## Abstract

Inversions have been proposed to facilitate local adaptation, by linking together locally coadapted alleles at different loci. Classic prior work addressing this question theoretically has considered the spread of inversions in “continent-island” models in which there is a unidirectional flow of maladapted migrants into the island population. In this setting, inversions are most likely to establish when selection is weak, because stronger local selection more effectively purges maladaptive alleles, thus lessening the advantage of inversions. Here, we show this finding only holds under limited conditions. We study the establishment of inversions in a “two-deme” model, which explicitly considers the dynamics of allele frequencies in both populations linked by bidirectional migration. For symmetric selection and migration, we find that stronger local selection increases the flow of maladaptive alleles and favours inversions, the opposite of the pattern seen in the asymmetric continent-island model. Furthermore, we show that the strength and symmetry of selection also change the likelihood that an inversion captures an adaptive haplotype in the first place. Considering the combined process of invasion and capture shows that inversions are most likely to be found when locally adaptive loci experience strong selection. In addition, inversions that establish in one deme also protect adaptive allele combinations in the other, leading to differentiation between demes. Stronger selection in either deme once again makes differentiation between populations more likely. In contrast, differentiation is less likely when migration rates are high because adaptive haplotypes become less common. Overall, this analysis of evolutionary dynamics across a structured population shows that established inversions are most likely to have captured strongly selected local adaptation alleles.

## Introduction

Chromosomal inversions are a form of structural variant that suppress recombination between loci. Inversions can result in reduced fitness due to the disruption of genes around their breakpoints (Kirkpatrick 2010), or from the capture and accumulation of deleterious alleles due to their lower effective recombination rate (Wasserman 1968; Berdan et al. 2021). Furthermore, inversion heterozygotes may experience reduced fecundity as a result of improper meiosis that results in aneuploid gametes (White 1978). Despite these negative fitness effects, the ubiquity of inversions has led to several putative explanations for their continued persistence (see reviews Kirkpatrick 2010; Wellenreuther and Bernatchez 2018; Faria, Johannesson, et al. 2019; Huang and Rieseberg 2020; Villoutreix et al. 2021). In particular, inversions could facilitate local adaptation under gene flow by increasing linkage between coadapted alleles and reducing effective migration of maladapted haplotypes (Kirkpatrick and Barton 2006).

Empirical evidence for this hypothesis has since been documented across a wide array of taxa (e.g. Lowry and Willis 2010; Cheng et al. 2012; Ayala, Guerrero, and Kirkpatrick 2013; Lee et al. 2017; Christmas et al. 2019; Faria, Chaube, et al. 2019; Huang, Andrew, et al. 2020; Koch et al. 2021; Hager et al. 2022; Harringmeyer and Hoekstra 2022), and a body of related theoretical work has also developed from the original model, investigating the roles of geography, chromosome type, and inversion length on the fate of adaptive inversions (Feder, Gejji, et al. 2011; Charlesworth and Barton 2018; Connallon, Olito, et al. 2018; Connallon and Olito 2021; Proulx and Teotónio 2022). For simplicity, this work often considers a “continent-island” model, in which inversions are introduced into an “island” population which receives maladapted migrants from a larger “continent” population. In this model, the selective advantage of an adaptive inversion is proportional to the rate of gene flow (Kirkpatrick and Barton 2006), and inversely proportional to the strength of selection on the island (Bürger and Akerman 2011; Charlesworth and Barton 2018). These results rely on the homogeneous maladaptation of migrant alleles which follows from the extreme migration asymmetry assumed between the continent and island populations (Kirkpatrick and Barton 2006). This scenario is unlikely to apply to many empirical systems, where local adaptation occurs in a structured population with greater symmetry and individuals migrate between similarly sized populations at rates that are similar to and from each population (e.g. Feder, Gejji, et al. 2011, Proulx and Teotónio 2022). With two-way dispersal, selection will interact with migration to determine the overall rate of maladaptive gene flow. However, there has been no thorough analytical dissection of the roles that migration and selection play individually in such a model.

In addition, it is important to consider not only whether an inversion spreads but also how the frequency of adaptive haplotypes affects their probability of being captured by an inversion. This has been briefly discussed before (Kirkpatrick and Barton 2006), and in relative terms when comparing X-linked and autosomal inversions (Connallon, Olito, et al. 2018). But so far models have sidestepped the problem by assuming that either an inversion capturing the coadapted haplotype simply existed or that such an inversion arose during a period of allopatry (Feder, Gejji, et al. 2011). Explicitly modelling the origin of the inversion is important because parameters favourable for the establishment of an adaptive inversion are not necessarily those where adaptive inversions are likely to arise. Assuming an inversion captures a random genotype, the probability of capturing a particular adaptive combination is proportional to its frequency. For example, adaptive inversions are expected to be favoured most when there are high rates of migrant gene flow, so there are fewer fit genotypes to be captured.

Here, we model the fate of locally adaptive chromosomal inversions in a two-locus, two-allele, two-deme model with migration and selection. We consider the case of symmetrical deme sizes and migration, as well as asymmetrical scenarios with the continent-island model as the extreme case. To understand the dynamics of inversions, we determine the probability of an adaptive inversion arising and its subsequent selective advantage in a population in which the locally adaptive alleles have reached their equilibrium frequencies and linkage under migration and selection. By considering the processes of inversion origin and spread in both demes, we determine population structures which favour the evolution of inversions that allow local adaptation under environmentally variable selection.

## Methods

We consider a population consisting of two demes linked by bidirectional migration with selection for local adaptation. We first derive analytical expressions for equilibrium allele frequencies at the local adaptation loci and the linkage disequilibrium (LD) between them. This will allow us to assess the frequency of each haplotype and hence the invasion probability of an inversion capturing a locally adapted combination of alleles. We then determine the probability of such an inversion arising and establishing itself in the population.

### Model

We model an infinite population of two demes, consisting of haploid, hermaphroditic individuals with discrete non-overlapping generations. The model is equally applicable to the case where there are two sexes at even sex ratio whose genetic determination is unlinked to the adaptive loci under consideration. Selection acts on two loci, *A* and *B*, that have two alleles each, *A_i_* and *B_i_*, where *i* ∈ {1,2} denotes the deme in which the allele provides a benefit *s_i_* (equal between the two loci). The relative fitness of an individual in deme *i* is either (1 + *s_i_*)^2^, (1 + *s_i_*) or 1, depending on whether it carries two, one or no allele(s) conferring local adaptation to its environment.

The life cycle begins with adults. These individuals reproduce, whereby pairs of parents are sampled according to their relative fitness in their current deme to produce one joint offspring. During reproduction, recombination occurs between the parental chromosomes (and their loci for local adaptation) at rate *r*. When alleles are held in an inversion, the recombination rate with non-inverted chromosomes drops to zero (double cross-overs and gene conversion are ignored). Migration between demes then occurs such that a proportion *m_kl_* of juveniles in deme *l* are migrants from deme *k*. After migration, the juveniles in each deme become the adults of the next generation. As the life cycle consist of just two phases, reproduction/selection and dispersal, the order of events within a generation does not affect the results.

At the beginning of a generation, *A_i_B_j_* adults in deme *k* are at proportion 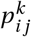 and have fitness 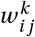. Among the parents sampled for reproduction, the frequencies are 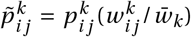, where 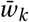 is the mean fitness in deme *k*. 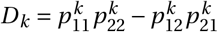 is the coefficient of linkage disequilibrium in deme *k*, and 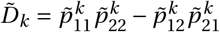 is the linkage disequilibrium after selection, among parents. Among the juveniles of the next generation, the frequency of genotype *A_i_B_j_* in deme *k* after migration, is given by

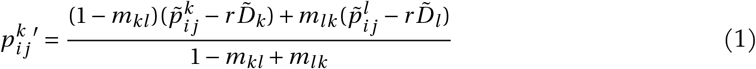

if *i* = *j*, and

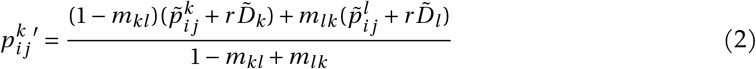

otherwise.

When migration is limited to one direction (i.e., *m*_12_ or *m*_21_ = 0) or when selection in one environment is very strong (*s_i_* ≫ *s_j_*), the model approaches the well studied “continent-island” model (hereafter superscript “C-I”, e.g., Kirkpatrick and Barton 2006 and Charlesworth and Barton 2018).

### Analysis

To use the quasi-linkage equilibrium (QLE) approximation, we first rewrite the genotype frequencies in terms of allele frequencies and LD, and then calculate their equilibria (Kirkpatrick, Johnson, and Barton 2002; Otto and Day 2011). This approximation assumes that recombination between the two loci is sufficiently high compared to migration and selection (*r* ≫ *m_ij_, s_k_*) to allow LD to reach an equilibrium much more quickly than the allele frequencies. This is justified here if we do not consider loci that are already tightly linked. But this is not an interesting case, because inversions then offer minimal advantage from suppressing recombination. To ensure the existence of an equilibrium, migration must also be weak compared to selection (i.e. max(*m*_12_, *m*_21_) < min(*s*_1_, *s*_2_)). These values allow the calculation of the equilibrium mean fitness in each deme, and hence the rate of increase of an adaptive inversion.

Using Equations 1 and 2 with *r* = 0, the dynamics of an *A*_1_*B*_1_ inversion are described by the transition matrix *M*_11_, in which the (*i, j*)-th entry describes an inverted adult experiencing selection in deme *i*, and whose offspring is located in deme *j* post-dispersal, given by

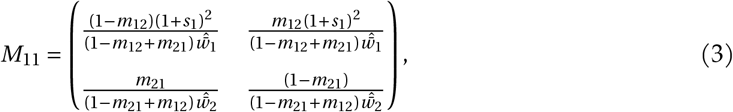

where 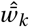 is the equilibrium mean fitness in deme *k* (we use the circumflex symbol ^ for equilibrium values throughout). The rate that a rare *A*_1_*B*_1_ inversion increases in frequency in the whole population (*λ*_11_) is given by the leading eigenvalue of *M*_11_. As the population is at equilibrium the growth rate of a recombining *A*_1_*B*_1_ haplotype is 1, so *λ*_11_ > 1 implies a benefit to the inversion that can be ascribed to the absence of recombination. From this measure of ‘invasion fitness’, we can approximate the invasion probability as 2(*λ*_11_ – 1) (Otto and Whitlock 2013). A similar transition matrix *M*_22_ can be derived for the behaviour of an *A*_2_*B*_2_ inversion (see File S1).

The invasion probability (2(*λ*_11_ – 1)) is specific to the *A*_1_*B*_1_ haplotype and hence conditional on an inversion capturing this allelic combination. To account for the probability of an inversion actually capturing the *A*_1_*B*_1_ haplotype in the first place, we need to take into account the frequency of this haplotype in a population at equilibrium. The simplest way of achieving this would be to multiply *λ*_11_ by the overall frequency of *A*_1_*B*_1_, across the two demes. This is an acceptable approach in the extreme case of the continent-island scenario, where the inversion is limited to the island and the growth rate only applies to that population. However, the overall frequency of *A*_1_*B*_1_ is no longer suitable in a two-deme model, because it gives equal weight to individuals in deme 1 where the haplotype is adaptive and and those in deme 2 where it is not adaptive. Accordingly, in order to determine the probability of an *A*_1_*B*_1_ inversion arising, we need to take into account not only the frequency of the *A*_1_*B*_1_ haplotype but also the relative reproductive value of the inversion in each deme. The reproductive value of the inversion in each deme is given by the left eigenvector of *M_ii_* and its components can be scaled to relative values that sum to 1. Call this scaled vector **v**_*i*_ = (*v*_*i*_1__, *v*_*i*_2__). Now, the probability that an inversion captures coadapted alleles (*A_i_B_i_*) and invades is given by

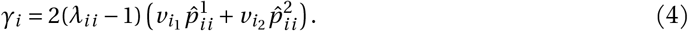

Finally, the probability of any locally adapted inversion establishing when it arises needs to consider both *A*_1_*B*_1_ and *A*_2_*B*_2_ haplotypes, and is given by

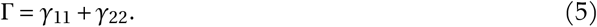

This is also equal to the probability of an inversion establishing itself overall, because inversions that capture allele combinations that are not advantageous in either deme (i.e. *A*_1_*B*_2_ or *A*_2_*B*_1_) are never favoured.

### Data availability

A Mathematica notebook containing derivations and code used to generate figures is available as File S1.

## Results

### Equilibrium allele frequencies and linkage disequilibrium

At equilibrium, the frequencies of the alleles (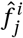 for allele *j* in deme *i*) are

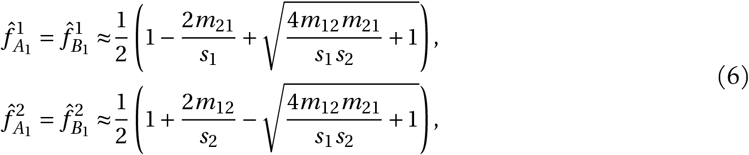

and the linkage disequilibrium between loci in deme 1 (*D*_1_) is

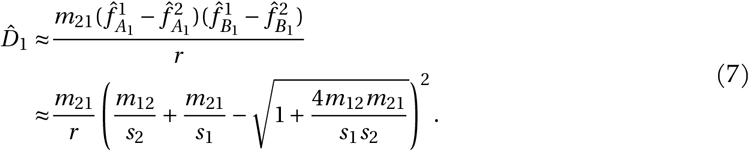

Linkage disequilibrium in deme 2 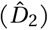 is given by replacing *m*_21_ with *m*_12_ and vice versa. These equilibrium values, derived here for haploidy and weak selection, are in accord with previous results (Akerman and Bürger 2014).

In the case where migration and selection are symmetric, *m_kl_* = *m* and *s_i_* = *s* (i.e., two populations with exactly opposing local selection pressures exchanging an equal proportion of migrants), the demes have symmetric allele frequencies 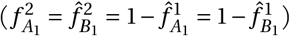 and linkage disequilibria (*D*_1_ = *D*_2_)

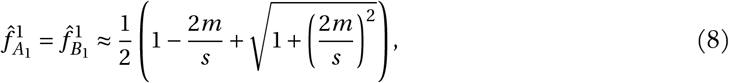

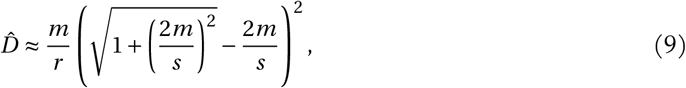

meaning that

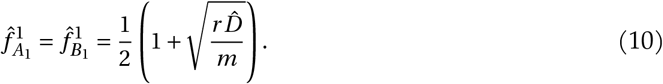

In the other extreme case, where there is unidirectional gene flow from deme 2 (“continent”) to deme 1 (“island”), the “continent” genotypes remain fixed to *A*_2_*B*_2_. Setting *s*_1_ = *s* and *m*_21_ = *m*

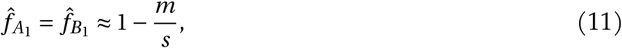

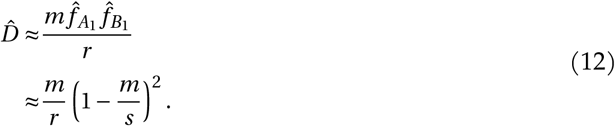

Locally adaptive alleles are more abundant in the symmetric scenario (equation 8) than in the continent-island scenario (equation 11). This difference arises because in the symmetric scenario a fraction of locally adapted migrants from a focal deme migrate to and survive in the other deme, only to return back and contribute to the frequency of beneficial alleles in the focal deme. In the continent-island scenario, in contrast, continental migrants can only introduce deleterious alleles into the focal deme.

In both scenarios, linkage disequilibrium is positive, indicating that the adaptive alleles tend to be found together in coadapted haplotypes (*A*_1_*B*_1_ and *A*_2_*B*_2_). This tendency increases with the strength of selection in both models 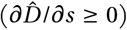, because selection favours the association of coadapted alleles, but decreases with the rate of recombination 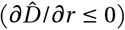 which breaks the coadapted haplotypes apart to create more intermediate haplotypes (*A*_1_*B*_2_ and *A*_2_*B*_1_).

The role of migration is less straightforward and differs between the two scenarios. At small migration rates, selection tends to be stronger relative to migration and demes are enriched for locally adapted haplotypes. Linkage disequilibrium then increases with *m* because more *A*_2_*B*_2_ combinations are introduced into deme 1 (and more *A*_1_*B*_1_ combinations are introduced into deme 2 in the symmetric scenario). When migration becomes higher, the balance between selection and migration shifts and migration tends to introduce proportionately more maladaptive haplotypes from the other deme, thus degrading the linkage disequilibrium that is built up locally by selection. The rate of migration at which this effect sets in depends on the model. In the continent-island scenario, migration decreases linkage disequilibrium when *m* > *s*/3. In the symmetric case, migration begins to decrease linkage disequilibrium at a lower point, when 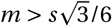, because the presence of *A*_1_*B*_1_ migrants in deme 2 generates more intermediate haplotypes through recombination. These individuals can back-migrate and degrade linkage disequilibrium in deme 1 (with the same process going on in the reverse direction).

### Invasion probability of a locally adaptive inversion

Having established the equilibrium composition of populations, we can now consider the fate of a new inversion that captures allele *A*_1_ and *B*_1_, which are locally adaptive in deme 1. We calculate the rate of increase and probability of fixation of this inversion. We again compare the two extreme models, the continent-island and the symmetric scenarios before examining the full model.

The growth rate of the inversion in the continent-island scenario is

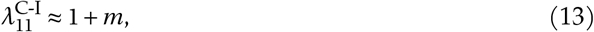

implying that migration is the main driver behind the selective advantage of inversions (Kirkpatrick and Barton 2006). The rate of growth is independent of the strength of selection in the island. The inversion’s benefit is the protection of locally adapted haplotypes from acquiring maladaptive migrant alleles through recombination. Increasing the strength of selection within the “island” has no effect to leading order, under the assumption that selection and migration are both weak.

In the symmetric scenario, the growth rate of an inversion is given by

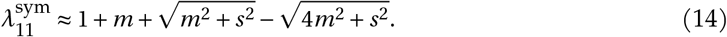

Since 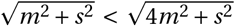, we always have 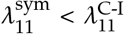 and the advantage of the inversion is weaker in the two-deme compared to the continent-island model. The inversion’s growth rate in the symmetric scenario now depends on the strength of local selection. Specifically, inversions are increasingly favoured with stronger selection (the square root terms in Equation 14 converge as *s* increases and 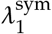 tends towards 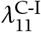). As the strength of selection increases, the proportion of deme 2 that is well-adapted increases. This means that new migrants carry more maladaptive alleles and recombination more often results in less fit offspring, so that the inverted haplotype has a greater advantage over non-inverted *A*_1_*B*_1_ haplotypes.

While stronger selection increases the frequency of maladaptive alleles among migrants, it will also remove them more effectively from the focal deme. This effect is not captured by our QLE approximation, so we numerically calculate the advantage of a rare inversion while assuming that allele frequencies are at the exact equilibrium calculated to second order in selection and migration (Figure 1). In the continent-island scenario, the genotypic composition of migrants is unaffected by selection. Stronger selection reduces the advantage of an inversion (as found by Bürger and Akerman 2011; Charlesworth and Barton 2018) because the island population becomes better adapted as selection increases, so that recombining adaptive haplotypes results in less fit offspring less often (Figure 1).

**Figure 1:**
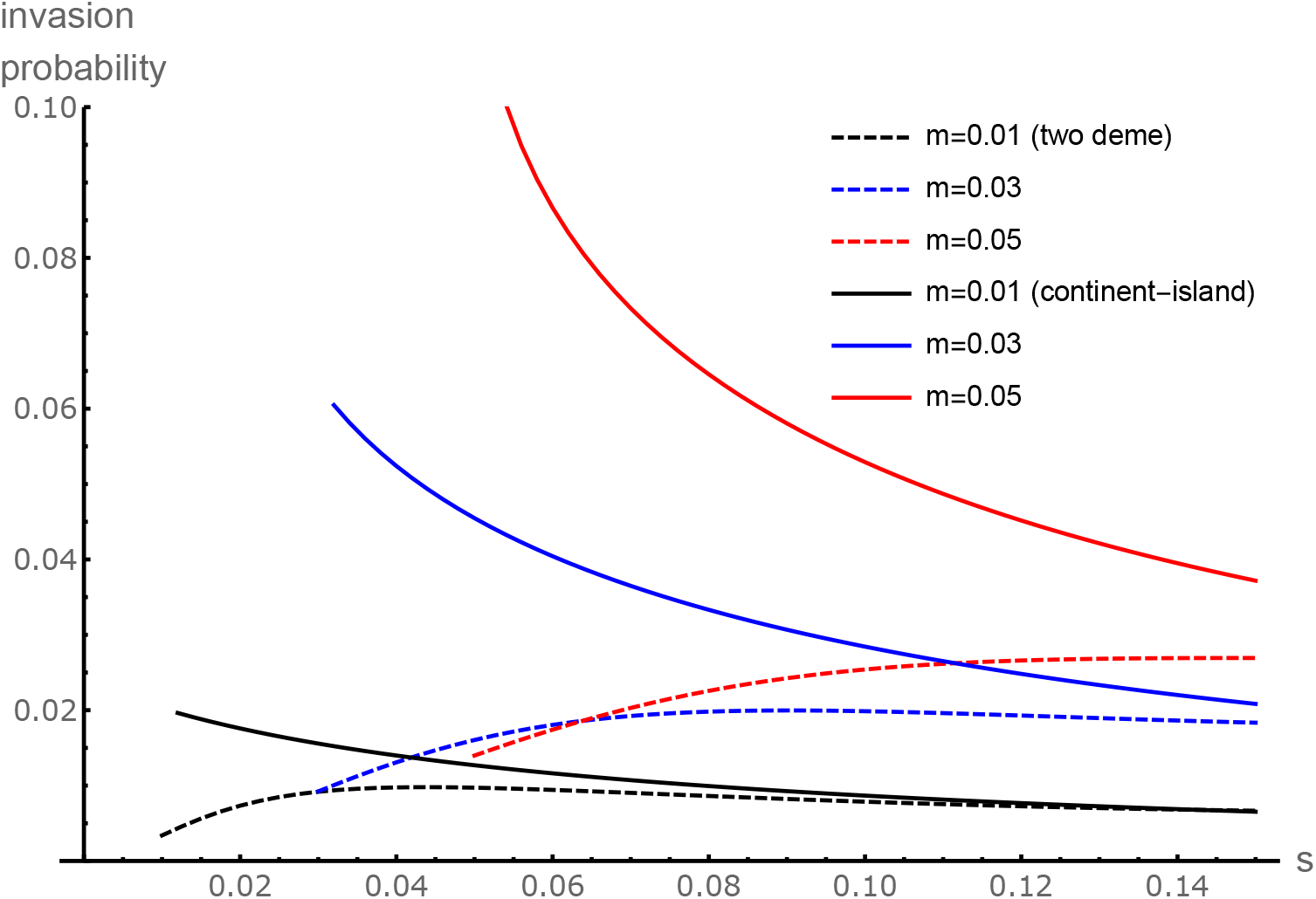
Invasion probabilities approximated to second order in migration and selection for an inversion capturing *A*_1_*B*_1_ in each of the symmetric and continent-island scenarios under various rates of migration.. Data with *s* < *m* are excluded as the adaptive alleles may not be at a stable equilibrium. The rate of recombination between the two loci was *r* = 0.15.

In the symmetric scenario, the numerical results confirm that increasingly strong selection favours inversions, as in the QLE results. This happens because selection reinforces local adaptation and makes migrants more maladapted. However, this advantage plateaus as the strength of selection increases, because adaptive alleles become more common. This decreases the advantage of inversions, as selection alone tends to weed out the maladapted combinations. Unless selection is very strong, the former force dominates, meaning that the selective advantage of inversions is primarily determined by the genotypic composition of migrants. Under very strong selection, the invasion probability under symmetric migration converges on that in the island-continent scenario (Figure 1), because the composition of migrants in each become similar.

Unlike the continent-island model, the two-deme model allows us to include asymmetric local selection and migration (Figure 2). Selection in the focal deme (*s*_1_) increases the degree of local adaptation and inversions therefore have a lesser advantage. This effect is strongest when there are more maladapted migrants entering deme 1 (higher *m*_21_, Figure 2A) or when the genotypic composition of migrants is more maladapted (higher *s*_2_, Figure 2C), but has a weaker effect on inversion invasion probability than parameters that change the genotypic composition of migrants (*m*_21_ and *s*_2_). For a fixed level of migration into deme 1 (*m*_21_), the growth rate of the inversion decreases with increasing migration out of deme 1 (*m*_12_) because inversions migrate out of the environment in which they are adapted (Figure 2B). Overall, a combination of increased migration from, and selection in, deme 2, are the most important factors in generating the inversion’s advantage (Figure 2D) — exactly the two parameters that are most extreme in the continent-island model.

**Figure 2:**
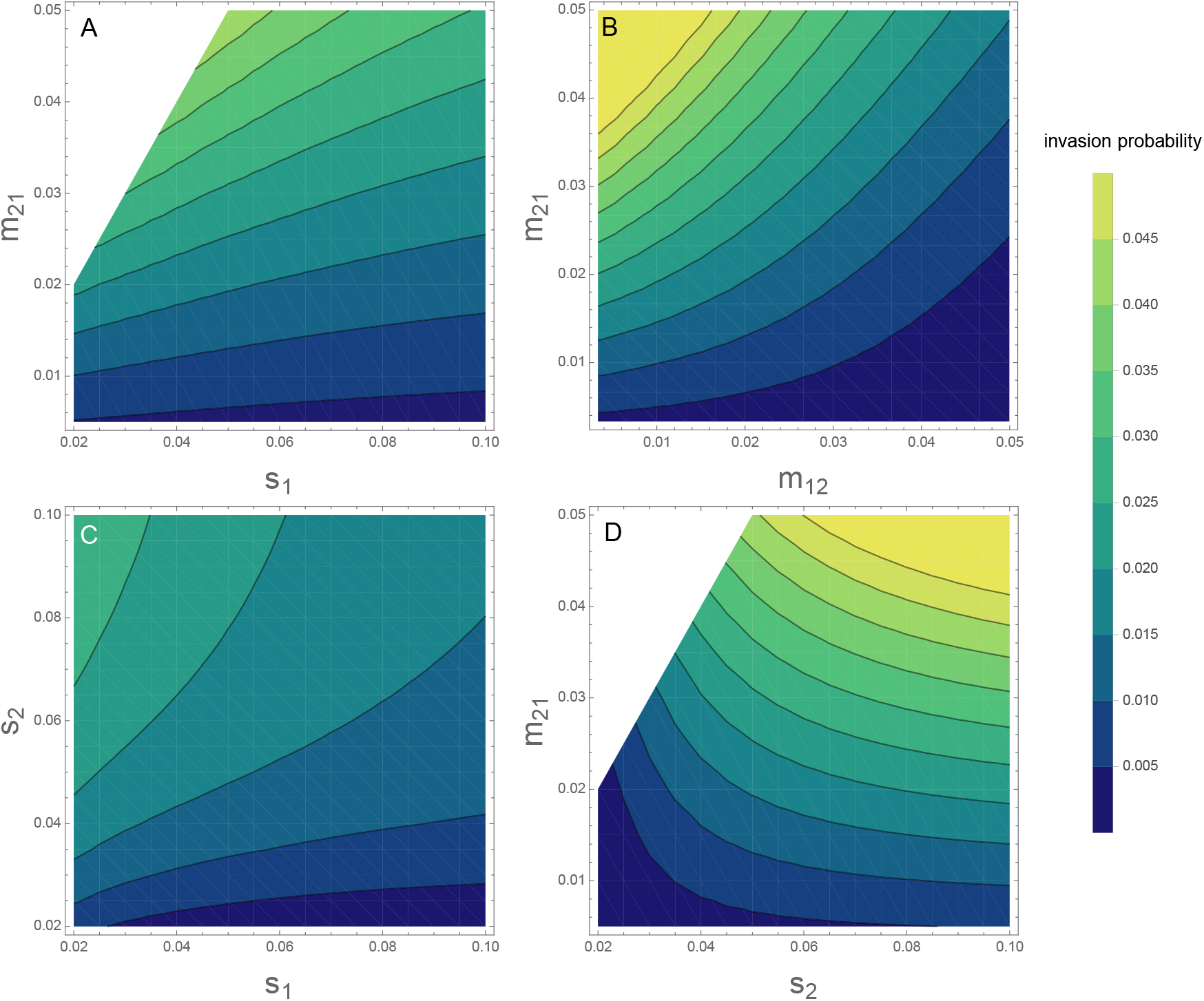
*A*_1_*B*_1_ inversion invasion probabilities calculated to second order in migration and selection terms. Where they do not vary, migration parameters are 0.02 and selection parameters are 0.05. Recombination was set to *r* = 0.15.

### Combined capture and invasion probability of locally adaptive inversions

The analysis above calculates the invasion probability assuming that an inversion captures the *A*_1_*B*_1_ haplotype. It does not take into account the probability that an inversion occurs in an *A*_1_*B*_1_ individual. It seems reasonable to assume that an inversion captures a random haplotype which means that the invasion probability should reflect the relative frequency of *A*_1_*B*_1_ as well as its reproductive value in each deme. Under this assumption, both the continent-island and two-deme scenarios predict similar patterns of invasion probabilities. As the strength of selection *s* increases, more locally adaptive genotypes are available to be captured by an inversion (Figure 3). The positive effect of selection on the frequency of locally adapted genotypes (*A*_1_*B*_1_) has a larger positive effect on the combined invasion probability than the negative effect of selection on the inversion’s subsequent selective advantage relative to the population (as illustrated in Figure 1). Thus, our results predict that stronger selection is more likely to drive the evolution of locally adaptive inversions. Importantly, this is true for both scenarions and radically alters the prediction for how inversions should contribute to local adaptation in the continent-island scenario (c.f. Figure 1).

**Figure 3:**
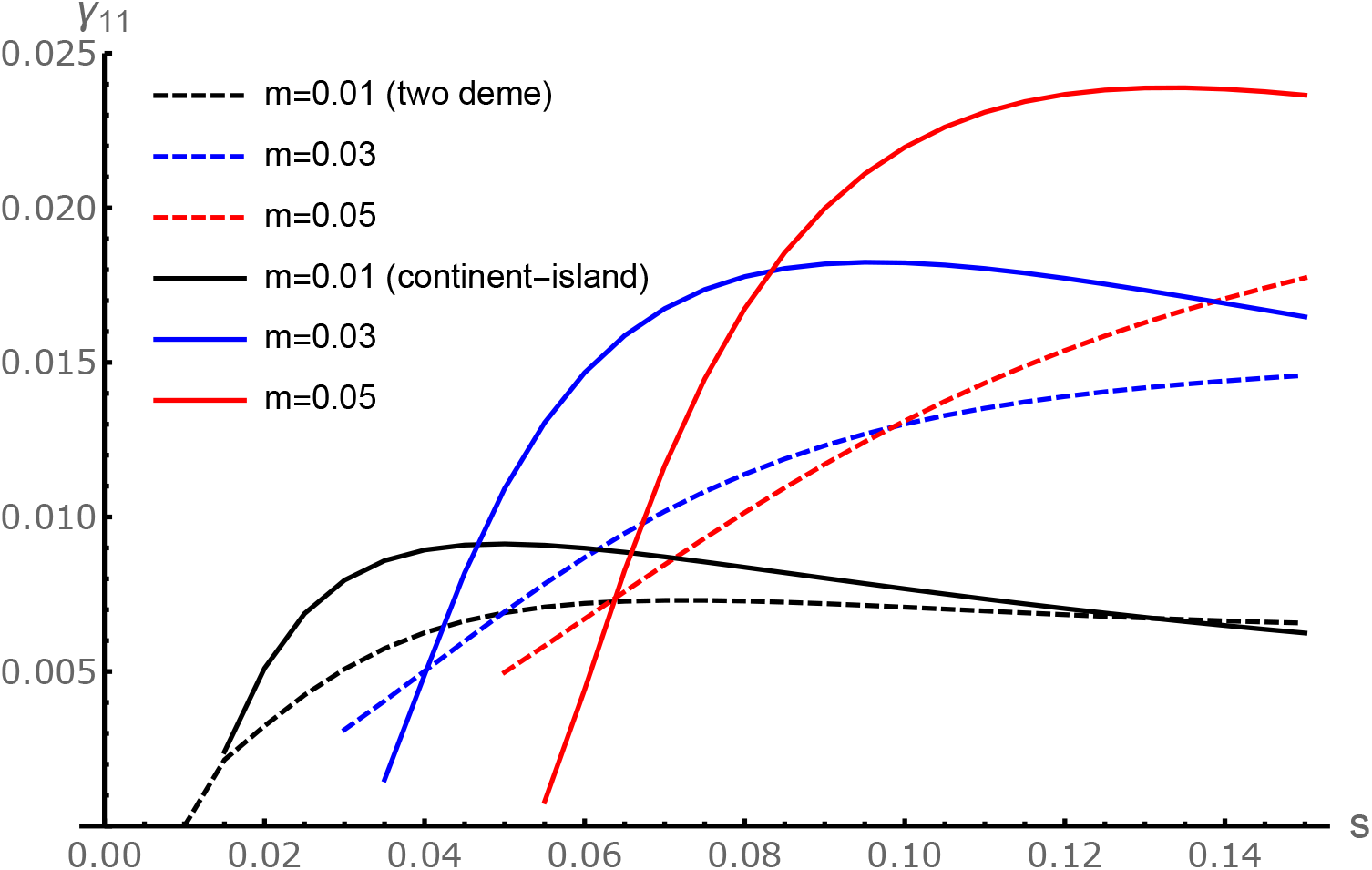
Combined probability of an inversion arising on an *A*_1_*B*_1_ haplotype and then invading (*γ*_11_). The invasion probabilities from Figure 1 are adjusted to account for the frequency and relative reproductive value of *A*_1_*B*_1_ in each deme. Equilibria are unstable for *m* < *s*, *r* = 0.15.

We can also see how asymmetric migration or selection affect the combined process of haplotype capture and invasion by inversions. While high migration into deme 1 strongly favours the invasion of existing adaptive inversions (Figure 2B), it also lowers the probability of them arising in the first place, due to the lower frequency of coadapted haplotypes. Thus, adaptive inversions are most likely to form and invade when *m*_21_ is intermediate, such that the probability of an inversion capturing an adaptive haplotype and the inversion’s subsequent selective advantage are both reasonably large (Figure 4A).

**Figure 4:**
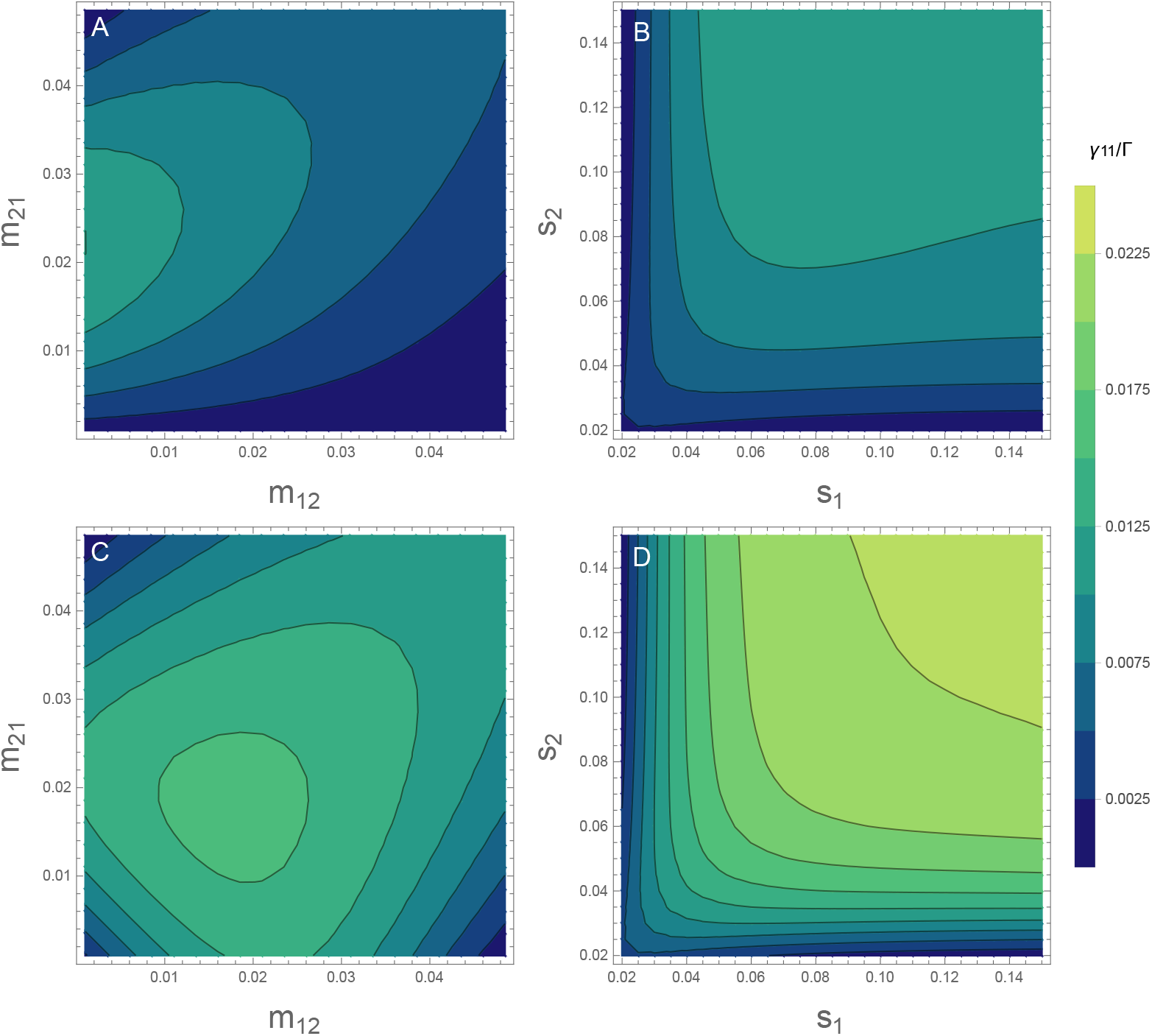
Total establishment probability of an adaptive inversion across the whole population. A, B: Combined probability of an inversion arising on the *A*_1_*B*_1_ haplotype and then invading (*γ*_11_) for asymmetric migration (A) or selection (B). C, D: Probability of an inversion capturing either adaptive haplotype (*A*_1_*B*_1_ or *A*_2_*B*_2_) and invading (Γ) for asymmetric migration (C) or selection (D). The continent-island model corresponds to *m*_12_ = 0 (Y axis in A, C) and the symmetric two-deme model corresponds to the *s*_1_ = *s*_2_ diagonal in panels B and D. Unless varying along axes, *m*_12_ = *m*_21_ = 0.02 and *s*_1_ = *s*_2_ = 0.05. To ensure stability, we vary parameters in the range where max(*m*_12_, *m*_21_) < min(*s*_1_*s*_2_), *r* = 0.15.

Increasing the strength of selection in either deme typically increases the chance that adaptive inversions will arise and spread. Increasing the strength of selection in deme 2 (*s*_2_) increases migration load and therefore the inversion’s advantage and increasing selection in deme 1 (*s*_1_) increases the probability of capturing the adaptive haplotype (Figure 4B). Yet, as discussed above, *A*_1_*B*_1_ inversion invasion probabilities decline under very strong selection in deme 1 (very high *s*_1_) by increasing preexisting adaptation. Nevertheless, stronger local selection usually creates a more favourable environment for adaptive inversions to arise and proliferate.

So far, we have only considered the evolution of a specific inversion, adaptive in one deme. This is the only plausible scenario in the continent-island scenario, where only inversions that capture the island-adapted haplotype *A*_1_*B*_1_ are of interest. However, with two demes, divergent local adaptation can occur from either adaptive inversion, both due to the beneficial effects in the favoured deme and due to the protection from deleterious recombination that such an inversion offers to individuals adapted to the other deme. So in this final section we consider the overall probability of local adaptation through the spread of an inversion that arises anywhere in the population (Γ:= *γ*_11_ + *γ*_22_; Figure 4C, 4D).

Under symmetric local selection, inversions are most likely to establish when migration is symmetric and intermediate (Figure 4C). Migration rates that are favourable for the establishment of inversion in one deme are not so favourable in the other (*γ*_22_ values can be seen by reflecting Figures 4A, 4B across the diagonal) such that symmetric migration rates give the highest overall probability of inversion establishment. Similarly, when migration is symmetric, strong and symmetric local selection is most conducive to the formation and spread of locally adaptive inversions (Figure 4D). Across both demes, this maximises the probability of capturing an adaptive haplotype while maintaining migration load.

## Discussion

Here, we have examined the evolution of locally adaptive chromosomal inversions while explicitly modelling selection across a structured population. Inversions can keep locally favoured allele combinations together in the face of maladapted migrants. Therefore, adaptive inversions spread fastest when migrant alleles are homogeneously maladaptive, as assumed in the continent-island scenario that has been well studied (Kirkpatrick and Barton 2006; Charlesworth and Barton 2018). The continent-island scenario represents an extreme, where migrants are fixed in their genetic composition, being purely maladaptive, with the migration rate alone determining selection for the inversion. In comparison, the two-deme model leads to a number of novel insights. By including the dynamics of selection and migration in the source population, we find that inversions capturing alleles experiencing relatively strong selection are more favoured, unlike the condition found when migration is unidirectional in the continent-island scenario (Figure 1). Extending the model to account for the probability that inversions initially capture favourable haplotypes shows that relatively strong selection is most likely to underlie inversions (Figure 3) and continent-island scenarios aren’t necessarily most conducive to inversion evolution (Figure 3). We further examine asymmetric selection pressures across demes, showing that strong selection in either deme generally promotes the establishment of adaptive inversions by either increasing the selective advantage or the probability of capture (Figure 4). Overall, our results suggest that inversions are particularly likely to arise and establish when selection on locally adaptive alleles is strong.

Theories concerning the origins of adaptive inversions can broadly be split into three categories (Schaal, Haller, and Lotterhos 2022): “capture”, in which an inversion creates a linkage group of existing adaptive variation and spreads (Kirkpatrick and Barton 2006); “gain”, in which an inversion is initially polymorphic (e.g. due to drift, underdominance, or acquisition of a good genetic background), and then accumulates adaptive variation which is subsequently protected from recombination (e.g. Lamichhaney et al. 2016, Samuk et al. 2017); or “generation”, in which adaptive variation is created when the inversion occurs through the breakpoint disrupting coding sequence or gene expression (Feder and Nosil 2009; Villoutreix et al. 2021, e.g. Jones et al. 2012). Our work focuses on the “capture” hypothesis in which locally adaptive alleles are already segregating and have reached migration-selection equilibrium and may have already evolved enhanced local fitness. This scenario is the most analytically tractable, and hence we analyse it here. However there is *a priori* no reason why any inversion with “capture” origins could not subsequently gain more adaptive variation at a later date as set-out in the “gain” hypothesis. In a pure “capture” scenario, we show large effect alleles are the most likely to underlie adaptive inversions.

The evolution of the effect size distribution of locally adaptive alleles is likely to favour those that are strongly selected, facilitating the evolution of adaptive inversions. In the short term, locally adaptive alleles must experience fairly strong selection to be able to resist being swamped by migration (Lenormand 2002; Yeaman 2015). Small effect alleles can still contribute to local adaptation when they arise in close linkage with large effect alleles, resulting in aggregated regions of adaptation which could be modelled as a single locus of large effect (Yeaman and Whitlock 2011). Alternatively, they can contribute transiently before being lost (Yeaman 2015). With high gene flow, and over long timescales, the architecture of local adaptation is expected to evolve towards a few, highly concentrated clusters of small effect alleles linked with large effect alleles (Yeaman and Whitlock 2011), which are likely to be particularly conducive to inversion establishment.

Migration regimes under which inversions are likely to form and spread are fairly specific because they must satisfy multiple requirements. Firstly, we assume that locally adaptive alleles are polymorphic, which means they must be able to resist swamping by migration. This condition requires relatively weak migration and is likely to be a significant constraint on the evolution of local adaptation (Feder, Gejji, et al. 2011). Then, given that locally adaptive alleles are maintained, higher migration rates favour the spread of inversions because they increase the frequency of the maladaptive alleles and thus the cost of recombination (Figures 1, 2). However, this also has the effect of reducing the frequency of adaptive haplotypes so that inversions are less likely to capture a full complement of adaptive alleles (Figure 4). The result is that higher migration rates do not always favour the evolution of inversions. In general, rates of migration may turn out to restrict the evolution of capture-origin inversions more than previously though.

Schaal, Haller, and Lotterhos 2022 used simulations to study the invasion of inversions capturing variation that influences a polygenic quantitative trait, finding that inversions involved in local adaptation tended to exhibit more of a capture than a gain effect when alleles were unlikely to be swamped. When alleles were prone to swamping by migration, persisting locally adaptive inversions had often gained much more adaptive variation post-capture. Under high rates of gene flow both capture and gain scenarios are plausible, depending on the effect size of the loci captured. Because adaptive alleles can be gained after the inversion arose and spread, recent inversions may offer the best opportunity to test our predictions about the effect size of alleles driving the evolution of locally adaptive inversions. The allelic content of such inversions could depend on how long the populations in question have been diverging, with the expectation that long periods of divergence results in a more concentrated architecture (Yeaman and Whitlock 2011). However, separating the individual trait effects of different loci within the inversion is challenging once they have been linked together. Thus, despite the prevalence of putatively adaptive inversions, mapping of quantitative trait loci has been achieved in only a handful of cases (e.g. Peichel and Marques 2017; Koch et al. 2021; and Poelstra et al. 2014 for an example unrelated to local adaptation) leaving open questions about the number and effect size of loci that underpin inversion selective advantage (Tigano and Friesen 2016).

We only consider the evolution of inversions that link alleles at two relatively nearby loci. It is possible that an inversion could capture more than two loci that affect local adaptation. As the number of loci contributing towards adaptation increases, it becomes less likely that an inversion will capture all the adaptive alleles on the same haplotype. Nevertheless, inversions will still spread if they capture more locally adaptive alleles than the population mean. A similar process has been proposed for the evolution of inversions that happen to capture fewer deleterious mutations than average (Nei, Kojima, and Schaffer 1967; Jay et al. 2022; Lenormand and Roze 2022). The relationship between invasion fitness and haplotype frequencies as the number of loci increases remains to be explored, but we expect inversion evolution will continue to depend on a balance between the selective advantage of the captured haplotype and on the probability of capturing a favourable haplotype.

Our model does not include deleterious mutations or breakpoint effects, which can affect the fate of inversions. Low rates of gene flux within inverted arrangements means that deleterious variation captured by the inversion persists for a long time throughout lineages, as purging this variation relies on rare events such as gene conversion and double crossover events. Inversion breakpoints can also disrupt gene function and result in lower individual fitness (White 1978; Kirkpatrick 2010), though this can occasionally be adaptive (e.g. Corbett-Detig 2016). These effects can be incorporated into the model by introducing a fixed cost or benefit. Reduced recombination within inversions severely weakens the efficacy of purifying selection on new mutations (Charlesworth 1996; Betancourt, Welch, and Charlesworth 2009). Mutation accumulation is particularly important while the inversion is at low frequency, because most inverted chromosomes will occur in heterokaryotypes where recombination is suppressed (Navarro, Barbadilla, and Ruiz 2000), though gene conversion and double crossover events may alleviate this a little (Berdan et al. 2021). We model a haploid population, but in diploids the presence and accumulation of strong recessive mutations within inversion will result in negative frequency-dependent selection which limits inversion frequency and the recombination rate (Nei, Kojima, and Schaffer 1967; Wasserman 1968; Ohta 1971). The generally deleterious effects associated with inversions likely mean that their invasion probabilities are much lower than we obtain here.

In summary, our results emphasise the likelihood that strongly selected loci can contribute to local adaptation in two ways: by increasing the frequency of adaptive haplotypes that can be captured by an inversion, and by increasing the rate of migrant gene flow and thus the potential cost of recombination. High migration rates also increase this recombination load and thus the selective advantage of an inversion, but this also reduces the frequency of adaptive haplotypes. The probability of adaptive inversion formation could be as important as its selective advantage in determining where such inversions are likely to be found.

